# Recurrent cortical networks encode natural sensory statistics via sequence filtering

**DOI:** 10.1101/2024.02.24.581890

**Authors:** Ciana E. Deveau, Zhishang Zhou, Paul K. LaFosse, Yanting Deng, Saghar Mirbagheri, Nicholas Steinmetz, Mark H. Histed

## Abstract

Recurrent neural networks can generate dynamics, but in sensory cortex it has been unclear if any dynamic processing is supported by the dense recurrent excitatory-excitatory network. Here we show a new role for recurrent connections in mouse visual cortex: they support powerful dynamical computations, but by filtering sequences of input instead of generating sequences. Using two-photon optogenetics, we measure neural responses to natural images and play them back, finding responses are boosted when inputs are played back during the correct movie dynamic context— when the preceding sequence corresponds to natural vision. This sequence selectivity depends on a network mechanism: earlier input patterns produce responses in other local neurons, which interact with later input patterns. We confirm this mechanism by designing sequences of inputs that are boosted or attenuated by the network. These data suggest recurrent cortical connections perform predictive processing, encoding the statistics of the natural world in input-output transformations.

## Introduction

A defining feature of the cerebral cortex is its extensive local recurrent connectivity. Excitatory cortical cells make up approximately 80% of the neurons in all cortical areas, and each excitatory cell receives hundreds or thousands of inputs.^1^ A majority of those input synapses come from other local neurons, within a distance of a few hundred microns.^2–4^

However, in many cortical areas, the function of this extensive local recurrent network is not well understood. For example, the sensory functions of primary visual cortex (V1) have often been explained by feedforward or feedback inputs alone, with little need for recurrent computations. Foundational work on V1 from Hubel and Wiesel^5,6^ showed that feedforward summation of visual thalamic inputs can create the edge detection properties (i.e. orientation and direction tuning) that are a defining characteristic of early visual cortex. Later work using inactivation and intracellular recording,^7–9^ confirmed this, finding that even though recurrent inputs to V1 neurons were large, recurrent input did not substantially refine visual tuning, for example not sharpening orientation responses.^10^

However, other work did find some contribution to visual function made by recurrent connections. Connectivity studies that combined *in vitro* and *in vivo* measurements showed neurons with similar orientation tuning are preferentially recurrently connected.^11–13^ This observation suggested that the recurrent network could perform preferential amplification of feedforward inputs driving orientation tuning^3,14^. That prediction was borne out using two-photon holographic stimulation, which confirmed that cells with similar orientation sensitivity did produce amplified responses when stimulated simultaneously.^15,16^

In sum, many years of study of V1 responses has found a primary role of the area is to extract edges from the visual field,^17,18^ and also found that the recurrent network of V1 facilitates that function.

Population-level recording studies confirm this idea, showing that much variation in V1 neurons’ responses is well-described by receptive field estimates derived from simple edges.^19,20^

But this framework cannot be the whole story. When V1 responses are recorded during natural vision, e.g. in response to natural movies, there is significant variation in neurons’ responses that are not well-explained by mere edge detection. That is, V1 response estimates derived from the tuning of the classical receptive field “center” do not come close to predicting all the variance of V1 neurons during natural vision. As Olshausen et al. put it,^21^ “What is the other 85% of V1 doing [during natural vision]?”. Up to 85% of the variability of V1 responses has not been explained by orientation or direction-based receptive field models, both when that question was posed and at the present time.

One possibility is that much of the responses of V1 neurons are explained by non-visual factors. Over the past 15 years, a large number of studies have focused on non-visual responses in V1 — responses influenced by the self-motion of the organism (e.g. running,^22–24^ top-down inputs,^25,26^ prediction,^27^ or cross-modal responses to non-visual sensory input, such as tactile sensation or auditory stimuli).^28,30^ but see^31^

On the other hand, it may be that the remaining unexplained V1 function is due to visual properties that have not yet been uncovered. There are hints that unknown visual responses drive this missing variance. The first hint of this sort is that V1 responses to natural movies are stronger and sparser — statistically quite different — than predicted from models that use V1 receptive field (edge) properties alone,^32–34^ implying some lawful rules of visual processing that have not yet been identified. Also suggesting there may be some V1 role in complex natural vision, neurons co-activated in natural vision are also more likely to be connected.^35^ And using large-scale recording methods, the responses of V1 neural populations have been found to be high-dimensional, as if they describe a very large space of visual responses extending beyond classical oriented-edge receptive fields.^36^ In fact, studies of V1 receptive fields generally observe a ‘non-classical surround’ — a large region outside the classical receptive field that affects V1 neurons’ responses, with hints of complexity^37^ that have not been so far fully deciphered (reviewed in ^38^).

One thing that could explain the missing variance in visual cortical responses is information about time. During natural vision, the visual input the organism receives changes as the visual world changes, both due to motion of objects and due to the organism itself moving. That means that the natural dynamic stimuli which produce strong and sparse V1 responses contain complex motion of various sorts.

But a major challenge to this idea — that the missing variance in V1 responses is explained by temporal information — is past studies have found only simple modulation of V1 neurons’ activity with time. Responses of V1 neurons to a sequence or movie of full-field stimuli are predicted by linear combinations of responses to single component stimuli,^19,39,40^ and V1 neurons do not have complex spatio-temporal receptive fields (STRFs) as have been seen in audition.^41–43^ That is, receptive field models that explain V1 responses as well as currently possible are generally spatio-temporally separable, meaning that the temporal kernel and spatial kernel of a V1 receptive field can be simply multiplied together to give its spatio-temporal response.^44^ ^34,45^

We thus are left with a major question about visual cortical responses and the V1 recurrent network. Much of what we can explain of V1 function can be explained by feedforward input and edge processing, but this does not explain V1 population responses to natural vision. What explains this missing variance in V1, and is that unknown function created by processing in the recurrent network?

Here we provide an answer to that question, showing that V1 responses are indeed modulated by temporal context, and that the recurrent network encodes features of natural vision, linking responses of neurons across time. We use two-photon holographic stimulation to replay patterns of input derived from natural movies. We find the gain, or magnitude, of the response to a given feedforward input is affected by other neurons’ activity in the recurrent network, showing that V1 responses to a visual stimulus vary in a predictable way based on temporal context. When a given patterned input is provided in the correct temporal context, preceded by the visual input that occurs in natural vision, the response to that patterned input is larger than when the same input pattern is delivered in a non-natural context. If one frame of a natural movie is preceded by the ‘correct’ prior input, as expected from natural vision, the response to that frame is boosted. Our data may explain why past studies have found a large ‘non-classical surround’ effect, but have not been previously able to find lawful rules of how simple stimuli predict non-classical surround responses: the non-classical surround is too complex to fully explain by exploring the space of visual stimuli using recording methods. Here, by replaying parts of known, identified natural stimuli using patterned stimulation, we show stimuli matching natural statistics specifically boost V1 responses, as populations of neurons in V1 responding to stimulus subparts influence each other through recurrent mechanisms.

Our results thus support the idea that the V1 recurrent network is performing complex computations over time not by generating sequential responses, but by filtering sequences of input. Natural vision produces a changing set of patterned input to the cortex, and either via experience or development, the cortical recurrent network has learned, and boosts, input sequences that correspond to natural vision. We confirm this, showing a network selecting natural sequences of input is consistent with our data by training an artificial recurrent network to boost certain sequences and showing that the network reproduces our experimental results. Then, returning to experiment, we use two-photon stimulation to identify a recurrent mechanism for this sequence filtering, construct novel sequences of input patterns following this mechanism, and show that sequences designed to be boosted do indeed produce larger responses. This confirms a recurrent origin for the sequence filtering that can underlie natural visual processing in V1.

Taken together, our results show that V1 neural responses are indeed sensitive to temporal context, but in a way that is not easily described by single cell response properties or receptive fields, but instead by populations of neurons affecting other populations via recurrent influences. Because we find that neurons’ gain is affected by the activity of other local neurons, the information for this boosting, or sequence amplification, process can be encoded in the large number of excitatory-excitatory synaptic weights in the V1 local network. These results thus provide one possible solution for what “the other 85%” of V1 is doing: beyond just edge extraction, the V1 recurrent network is sensitive to the complex temporal structure of the natural world.

## Results

Two-photon holographic stimulation (Fig. 1)^46,47^ allows us to deliver direct inputs to populations of cells in the cortex and directly study the influences of local neural activation on other local neurons. We use a viral expression strategy^48^ that co-expresses an opsin and a calcium indicator in excitatory neurons in layer 2/3 of mouse primary visual cortex, stably over days to weeks, with minimal optical crosstalk between imaging laser and opsin.^48^ This approach delivers optogenetic inputs that are precise spatially (Fig. 1F) and temporally, with one input pattern changing to the next within tens of milliseconds (Fig. 1I; 30 ms pattern duration).

**Figure 1:**
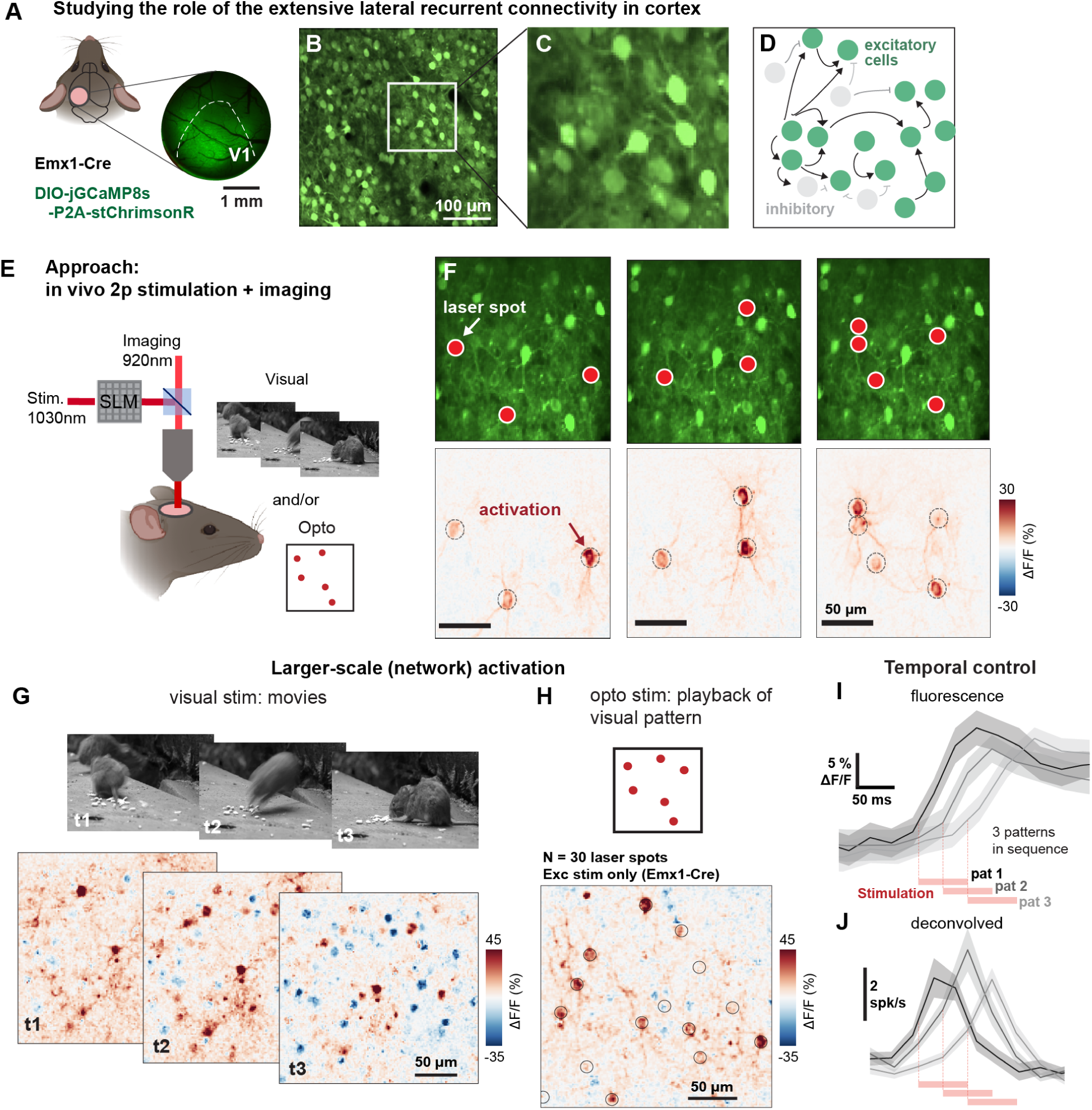
Using two-photon holographic stimulation and imaging to probe the role of excitatory recurrent connections. **(A)** 3mm window surgical implant over V1. **(B)** FOV image of excitatory cells expressing both GCaMP8s and opsin stChrimsonR (via AAV-DIO-jGCaMP8s-P2A-stChrimsonR in Emx1-Cre line). **(C)** Enlarged region of (b). **(D)** Schematic of the ubiquitous recurrent connections between excitatory cortical neurons. **(E)** Schematic of two-photon imaging and stimulation experiments, allowing pairing of visual and patterned optogenetic stimuli. **(F)** Example of spatial precision of 2p stimulation; left to right: three different patterns, with 3, 3, and 5 laser spots, each spot targeted to a neuron. **(G)** Network activation from natural movie input at three timepoints. **(H)** Network activation, with non-target changes in non-stimulated neurons, also results from optogenetic patterns (black circles: laser spots, 30 total, a subset of image shown here for visual clarity). **(I-J)**, Example of temporal precision of sequential 2p stimulation: 3 patterns, 15 cells each, 60 ms stim per pattern with 30 ms between pattern onset. **(I)** fluorescence traces, reflecting calcium responses of stimulated cells; **(J)** corresponding deconvolved traces using OASIS (Methods). These data show that even with some overlap, imaging can resolve distinct temporal peaks with stimulation onsets separated by 30 ms.

Two-photon optogenetics can mirror features of natural visual input by driving population activity in the cortical network. When animals are shown natural visual movies, many neurons in the visual cortex respond (Fig. 1G). Using two-photon stimulation we can simulate these activity patterns by giving input to many neurons at once (Fig. 1H; in this work we target 15-30 cells). This all-optical approach allows us to measure sensory responses, select neurons that respond, and replay activity patterns.

### Sequential order affects V1 neurons’ responses through non-target suppression

If recurrent effects influence V1 responses to temporally-varying input, then changing the order of patterns of input might be expected to produce differences in neurons’ responses. To determine if this is true, we chose patterns of neurons at random (three patterns; A, B, and C, each a fixed duration, 30 or 60 ms in different experiments, spanning a range of the temporal frequencies often seen in vision: e.g. frame times used in videos,^49^ flicker fusion frequencies,^50,51^ and photoreceptor dynamics.^52^ We presented these patterns in two different orders (ABC or CBA, Fig. 2A).

**Figure 2:**
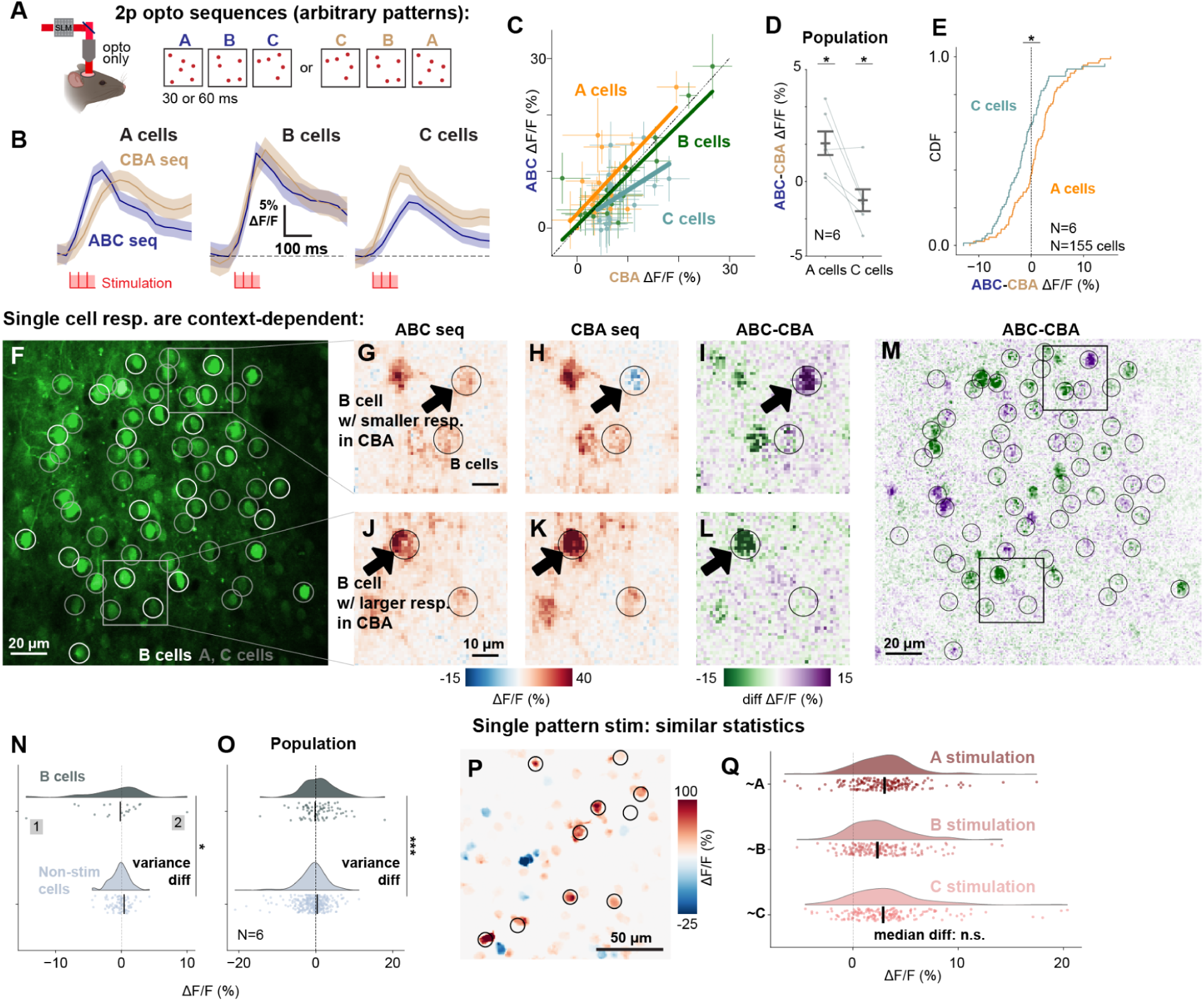
Patterns of input produce different responses in the V1 network depending on the sequential context. **(A)** Experiment schematic: stimulation of three distinct patterns (e.g. N=30 cells per pattern, cells chosen at random for each pattern). Two sequences: forward (ABC) and reverse order (CBA). **(B)** Responses depend on sequential order. Mean responses averaged across cells of A (left), B (middle), and C (right) cells in the ABC (blue) or CBA (beige) trials (example expt, pattern duration: 30 ms). **(C)** Points: average response (across trials) of individual neurons, same data as in B. Change in slope for A, B, C indicates differences in response based on sequential order. **(D)** Population data (N=6 expts). gray lines: ABC - CBA responses of A (left) and C (right) cells; averaged over A, C cells per experiment. Error bars: SEM. (t-test, A cells (N=79): p<0.0001, C cells (N=76): p=0.001). **(E)** ABC and CBA responses differ for A cells (N=79) and C cells (N=76) (K-S test, p<0.001). **(F-M)** B cell responses reveal that effects of sequential context vary from cell to cell. **(F)** Anatomical image (green: bicistronic GCaMP8s and stChrimsonR expression), white circles: cells stimulated in B pattern, gray circles: A and C pattern cells. **(G-L)** Neurons’ (B cells, black circles) responses to stimulation change based on which other neurons (A, C) are stimulated before them. B cells. **(G-I)**: example B cell with stronger activation in the ABC sequence compared to CBA. **(J-L)**: example B cell with stronger CBA activation. **(M)** Larger view of cells shown in g-i. Image matches anatomical FOV in f. Black squares: regions shown in g-i. **(N)** Some stimulated cells firing is increased and some decreased by previous stimulation pattern. x-axis: B cell response difference between ABC and CBA pattern; if responses were not modulated by sequence, variability would be similar to non-stimulated neurons (bottom, gray); instead it is larger (A-D test for variance diff, N=19 B cells, N=95 non-stim cells, expt shown in f-j, p<0.01). **(O)** Same as k, across N=6 expts (A-D test, N=81 B cells, N=339 non-stim cells, p<0.001). **(P-Q)** Sequence modulation is not due to response differences to single pattern stim: A,B,C patterns have similar statistics (data from b-e; Mood’s median test, N=3 stim, p=0.15).

We find the sequential order of these population input patterns does indeed affect neurons’ responses (Fig. 2B-E). The first pattern produces a greater response than the same pattern when it is delivered later in the sequence (Fig. 2B-E). This sequential context dependence occurs in the cells stimulated as part of the A, B, or C patterns. On average, the cells stimulated first in the sequence show stronger responses (Fig. 2C-E; the median of the A and C distributions differ in Fig. 2E). Larger effects are seen in individual neurons, which are strongly modulated by sequential position (e.g. there is significant variance in the distributions in Fig. 2E: 37% of A neurons and 45% of C neurons are significantly different from zero, K-S test p < 0.05).

To understand whether prior patterns of input to other cells could change the response of neurons to a particular input, we analyzed responses in the B cells (Fig. 2F). The B input pattern is preceded by one pattern in each sequential order: the A pattern in the ABC sequence, and the C pattern in the reverse sequence. Thus, if there is a specific effect of sequential context on individual neurons, the firing of some B cells to the B input should be modulated also by the pattern that precedes it. Indeed, we found that many of these cells were significantly modulated by sequential context (Fig. 2G-L). Even though the B input was always the same, some B cell responses were larger in the ABC sequence (Fig. 2G-I) and some were larger in the CBA sequence, as the pattern that preceded the B pattern was changed (Fig. 2J-L; variance differences via K-S test, Fig. 2N-O).

In sum, stimulation with sequences of random population input patterns shows that V1 neurons’ responses to fixed inputs are highly dependent on sequential context. Responses to an input pattern are affected by prior patterns of input to different neurons. This suggests that the visual cortex recurrent network may perform a time-based computation for sensory stimuli, and to understand this, we next investigated how sequential modulation relates to natural vision.

### Patterns corresponding to natural visual inputs produce larger responses

To study sequential modulation in the visual context, we turned to dynamic natural visual inputs. One of the most common kinds of sequential input that the visual cortex receives comes in response to natural motion. Because of the size and scatter of cortical receptive fields,^53,54^,and because axons carrying inputs contact many cells, the cortical network receives a changing set of patterned inputs during dynamic vision. As an example, when we see a person or animal running, the cortex “sees” a sequence of patterns of input.

In principle, the cortical network could store, in its recurrent connections, information about these patterns and thus the temporal structure of the natural visual world. This could allow the network to preferentially respond to sequences of input corresponding to natural vision. In that case, naturally-occurring sequences would be expected to produce different responses in the network than sequences that do not correspond to natural vision. This could occur by earlier parts of a dynamic natural stimulus influencing responses to input arriving at later times, as we observed with random input.

We examined this by using two-photon stimulation to mimic responses from one part of a natural movie while we changed the prior visual context. We first measured responses to a single frame taken from a movie (Fig. 3A). We constructed a two-photon input pattern from the cells activated by that single frame. We then showed animals the movie and played back the response to the frame with two-photon stimulation, replacing the visual frame with the input pattern either at the correct time in the movie (Fig. 3A,B: matched context), or at a different time, when the preceding movie frames were not matched to the stimulation pattern (Fig. 3A: unmatched context).

**Figure 3:**
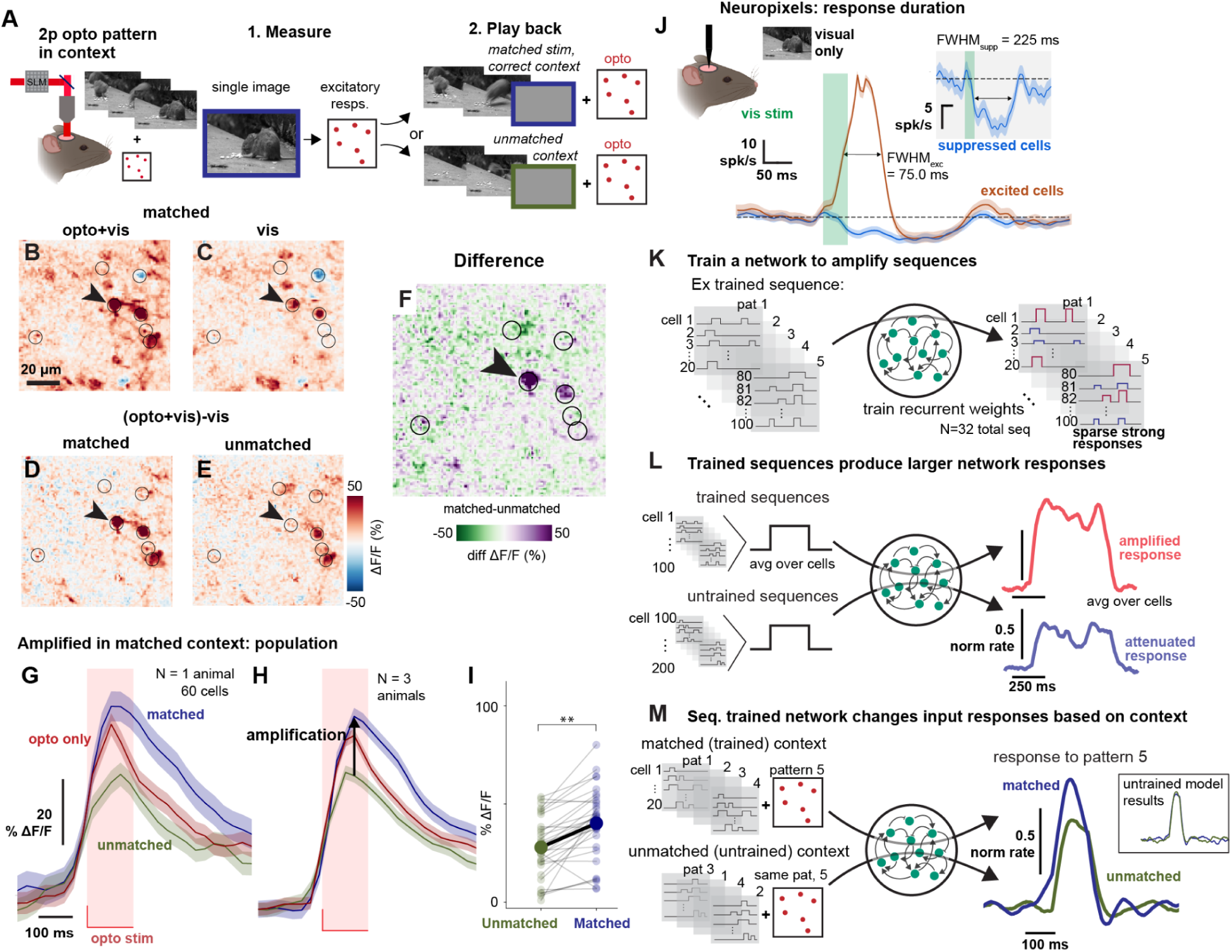
Sequence amplification for visual input, observed by playing back responses to a single frame of a movie. **(A)** Schematic of dynamic context playback experiment: We recorded responses to a frame from a natural movie, and played back that pattern using stimulation during a visual movie. We did this at the correct time in the movie (matched context) or an incorrect time (unmatched). **(B-C)** Example cell responses to combined opto and visual stimulation (B) and visual stim alone (C). **(D-E)** Responses in matched and unmatched contexts, computed by subtracting vis response from opto+vis combined response. Black circles: optogenetic stimulation targets. **(F)** Example larger response in matched context (arrow), difference of responses shown in (D) and (E). **(G)** Mean responses averaged across cells in matched (blue), unmatched (green), and opto only (red) trials (N=1 expt, pattern duration: 120ms). **(H)** Population time courses, N=3 animals, same conventions as (C). Responses are larger in matched context. **(I)** Points are cell responses in matched (blue) and unmatched (green) contexts from data in h. Mean optogenetic responses are significantly larger in the matched context (t-test, N=32 cells, p=0.004). **(J)** Electrophysiological responses to a flashed natural image show responses sustained for up to hundreds of milliseconds. Excited cells (N=967 cells, orange; errorbar, SEM, largely hidden by mean). Suppressed cells (N=342 cells, blue). FWHM: Full width at half max. **(K-M)** RNN model trained to amplify specific sequences shows responses that match our data. **(K)** Network is trained to amplify sequences of input, resulting in **(L)** Boosted output timecourses for trained input sequences. **(M)** Simulating experiment of (A-I) recapitulates the data: single patterns taken from a natural sequence produce larger responses in the correct (blue) context. Inset: untrained model does not show context-dependent responses. Error bars: SEM.

If V1 responses were dependent on the dynamic context of natural vision, neurons should produce different responses to the same, fixed input pattern when presented in different contexts. That is what we found (Fig. 3D-I). The input pattern produced a larger response in the sequential context matched to the visual movie than when presented in the unmatched context. Responses were not just increased in the matched context, but were attenuated in the unmatched context, relative to the response when it was preceded by an unchanging gray screen (Fig. 3G-I; Supp Fig. 2A).

An immediate question is about which mechanism can allow responses to one pattern of input to influence responses to later input. For a recurrent network interaction to modify responses to later inputs, it would seem that responses from one frame should be sustained somewhat in time. Then, one input pattern could produce non-target responses in other local neurons, and a later input could fall on these non-target neurons while their responses were changed by the previous input, allowing the two inputs to interact with each other. To determine if this kind of sustained response was present, we turned to Neuropixels electrophysiological recordings, due to their fidelity in measuring the times of spikes. We recorded V1 neuron responses to flashed single image frames (as in Fig. 3A) and measured the duration of their non-target responses. We sorted neurons into groups based on whether they showed activated or suppressed responses, and found that both activated and suppressed responses were sustained after visual stimulus offset, with suppressed responses lasting for several hundred milliseconds (Fig. 3J). This is sufficient for one frame of a natural visual movie to interact with the next. For comparison, a movie at 25 frames per second, a rate comfortably within the range where humans see smooth motion, has a 40 ms frame time. The mouse visual system likely supports even faster processing than the human, and yet here we found that natural images flashed for short times produced responses 75 ms long for activated neurons (Fig. 3J) and 225 ms for suppressed cells (Fig. 3J, inset). These data suggest that the dynamics of input during natural vision is sufficient to allow overlap between patterns, allowing recurrent influences from earlier input patterns to affect responses to later inputs.

### A recurrent neural network model that boosts specific sequences produces the context-dependent responses we see in our data

The observation that responses to single input patterns are modulated by context — by earlier patterns or frames (Fig. 3A-I) — implies that longer sequences of input are also selectively filtered, as each movie frame response interacts with later movie frames. That is, the pairwise sequential interactions we found, where single frames of input are boosted based on prior input, can also enable processing of longer sequences of dynamic input where longer natural sequences of input are boosted. Biological motion, like observing a predator moving to attack, can be ongoing for some time and generate changing sequences of input patterns to the cortex through that time.

To determine whether boosting of even longer sequences can result from the temporal interactions we studied, we examined a recurrent neural network (RNN) trained to preferentially generate stronger responses for some seconds-long input sequences and not others (Fig. 3K-M). We asked if it also showed the context-dependent effects seen in our experiment and found that it did. Indeed, when an RNN was trained to boost a sequence of input, a single pattern extracted from the sequence and played back in the correct context produced a larger response, compared to when it was presented in the incorrect context (Fig. 3M).

Thus, an RNN trained to boost a sequence of input (Fig. 3M) mirrors the context-dependent stimulation effect seen in our experiments (Fig. 3H). Together with the finding that neurons’ responses to input are dependent on earlier inputs (Fig. 2), these observations support the idea that the visual cortex recurrent network is filtering extended sequences of input, specifically boosting input sequences corresponding to natural vision.

### Recurrent network mechanism: one input pattern creates responses in other, non-targeted local neurons, which interact with later input patterns

If sequence modulation was created by local, recurrent interactions, we should be able to see signs of these interactions in the network activity. To look for specific influences — to see how particular input patterns filter later patterns — we separated stimulation patterns in time, inserting delays of several seconds (8 seconds, Methods) between one or more patterns in a sequence (Fig. 4A). We first inserted a delay between pattern B and C in an ABC sequence (A,B,C patterns randomly chosen, Methods).

**Figure 4:**
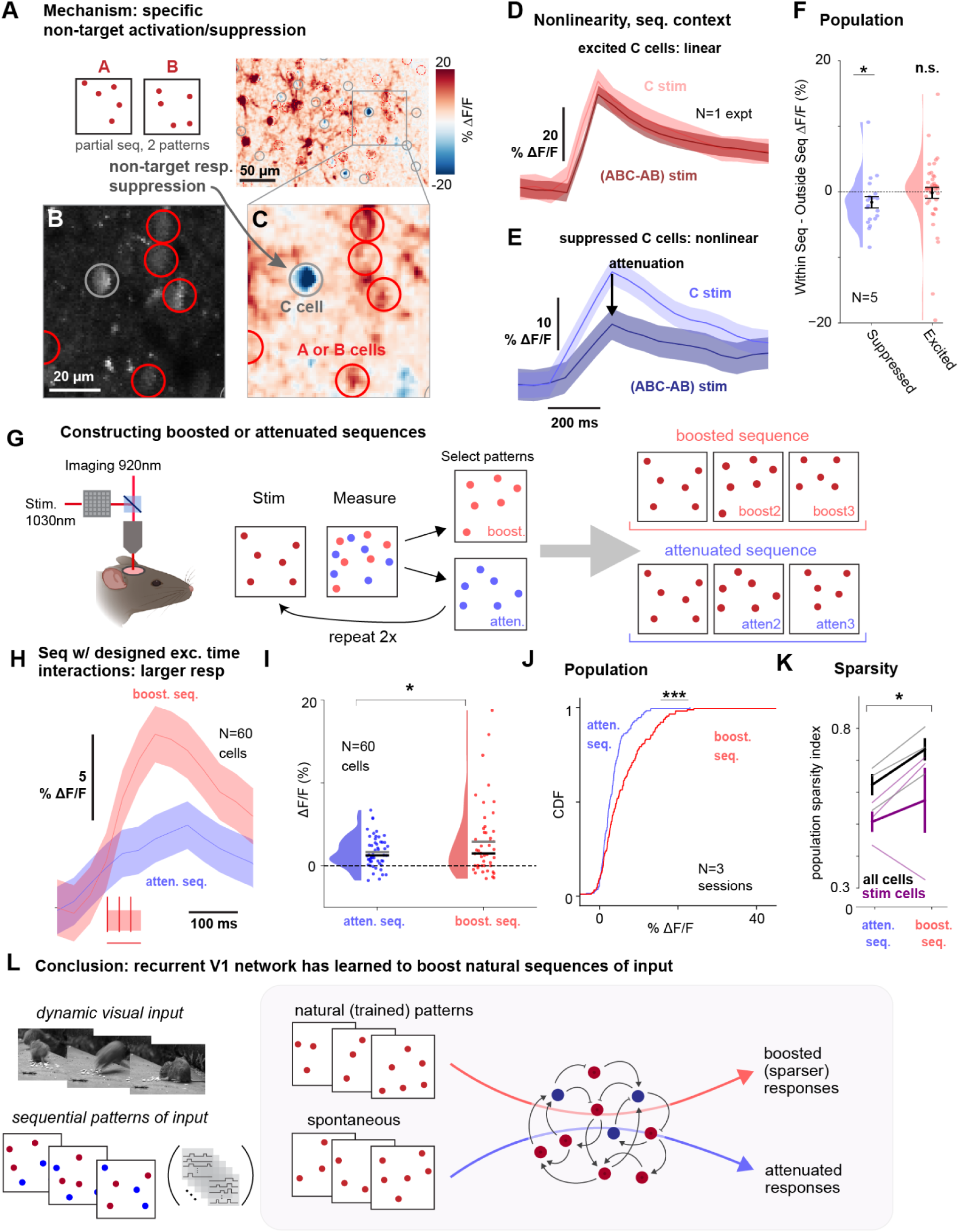
Sequences designed to be boosted produce strong and sparse responses, validating a recurrent network mechanism. **(A-C)** Separating A, B, and C patterns in time shows non-target responses induced by earlier patterns. Red circles: stimulated cells (from A or B pattern). Gray circle: C cells, here unstimulated. **(A)** Responses to AB sequence. Left, schematic; right, GCaMP responses. **(B)** Anatomical image (FOV; averaged over time). **(C)** Zoomed view shows suppression in unstimulated neuron (gray circle). **(D-F)** Excited non-target cells show linear summation, but suppressed cells show sublinear summation. **(D)** Excited cells from one example experiment (N=5 cells). Light red: response to C stimulus alone. Dark red: prediction, response to ABC sequence minus response to AB sequence. Response and prediction overlap. **(E)** Suppressed cells, with sublinear summation (N=5 cells; same expt). Same conventions as d. Response to C within the ABC sequence (ABC-AB, dark blue) is smaller than resp. to C alone (light blue; t-test, 300 ms period after stim onset, p=0.013). **(F)** Population data (N=5 expts; N=62 cells, C cells only; expt compares C stim, ABC stim, and AB+C stim as described in d-e). Blue: suppressed cells (response to AB stim<0; t-test vs linear prediction, N=20 cells, p=0.04). Red: excited cells (response to AB stim>0; t-test, N=42 cells, p=0.86). **(G)** Designing sequences to be boosted or attenuated based on non-target interaction mechanism: experimental schematic. We image responses to stimulation, and then select cells based on response to form the next pattern in the sequence. **(H)** Mean responses of boosted (pink) and attenuated (blue) sequences (N=20 cells per pattern, 3 patterns in the sequence; 30 ms pattern duration). **(I)** Distribution of responses of stimulated cells. Mean (gray horizontal line), median (black), means are sig. different (t-test, p=0.033; N=60 cells from N=3 patterns). **(J)** Population data (K-S test, p<0.0001, N=189 exc cells, N=179 supp cells, N=3 expts). **(K)** Sparsity is greater for boosted patterns. Each point: average across neurons. Sparsity in entire population is significantly greater in boosted sequences (t-test, p=0.011; N=3 expts; all cells, gray). **(L)** Mechanism and conceptual model from this work: the recurrent network filters sequential inputs, boosting some and attenuating others. Errorbars: SEM.

The reduced sequence produces striking non-target responses (Fig. 4A-C) in cells not receiving stimulation, with some non-stimulated cells activated, and others suppressed. These non-target neural responses are likely to be primarily due to local recurrent interactions, not due to axonal or dendrite activation, as here we used a somatically-targeted opsin (stChrimsonR).^48^

These non-target responses induced by the AB sequence can occur in neurons that will later be stimulated in the C pattern. This could be a mechanism for sequential modulation: that is, a single input pattern produces non-target responses in other local neurons, and these responses interact with later patterns of input to modify responses to those later patterns.

To see how earlier patterns impacted later patterns, we examined how C cells changed their responses when preceded by the AB pattern, and whether this change was predictable from the AB pattern responses. In prior work we have found attenuation-by-suppression,^55^ where V1 neurons, when suppressed, produce smaller (sublinear) responses to input. Such a nonlinearity could result in preferential suppression of some sequences, resulting in relative boosting of other sequences. To understand if sequence selectivity could depend on such nonlinearities, we examined non-target responses in C cells generated by the AB sequence. We compared the ABC response to a prediction computed by summing the C response and the AB response. We found that excited non-target cells did produce linear responses when they were stimulated next in sequence, with ABC responses well-predicted by the sum of non-target responses to AB and C (Fig. 4D,F, Supp. Fig. 3A). However, suppressed non-target cells showed nonlinear responses (Fig. 4E,F, Supp. Fig. 3A).

These responses suggest that the sequential modulation we observed does involve some network mechanisms. Non-target responses, induced by earlier patterns of input through the local recurrent network, generate a non-target response pattern that interacts with later patterns.

### Designed input sequences produce predicted effects, confirming the network mechanism

To provide an explicit test of this network mechanism, we constructed sequences that should be treated differently according to the mechanism described above.

To construct such sequences, we first measured responses to a single input pattern (Fig. 4G). We then created later stimulation patterns based on the non-target responses to those single patterns. We created ‘boosted’ sequences by adding a pattern of stimulated cells that were excited by the first pattern, and ‘attenuated’ sequences by adding a pattern of stimulated cells suppressed by the first pattern (Supp. Fig. 3G-J). We repeated this step a second time to create boosted and attenuated sequences that were three patterns long.

If our constructed sequences, when delivered to the network, produce the expected excitation or suppression, this would be strong evidence that this recurrent mechanism underlies sequence modulation.

We studied this by playing back these sequences onto the cortical population. Confirming our hypothesis, we found that the sequence designed to be boosted produced a larger response than the sequence designed to be attenuated (Fig. 4H-J). This interaction between non-target responses induced by one pattern affecting later patterns is therefore a mechanism where some sequences can be enhanced and others suppressed.

However, one additional nonlinear phenomenon resulted beyond the suppression of some sequences (Fig. 4A-E). Patterns designed to be boosted produced responses that were sparser than suppressed sequences — a number of outlier cells produced large responses that elevated the mean (Fig. 4I,K). This is a kind of nonlinear amplification effect produced by later patterns falling on non-target cells that were excited by earlier patterns. This sparsity is a hallmark of responses to natural movies,^32,33,56^ and supports the idea that the cortical network can selectively boost sequences of responses associated with natural vision. Thus we observe at least two kinds of nonlinearities that can contribute to sequence filtering, boosting some sequences and suppressing others: first, the attenuation-by-suppression effect that suppresses patterns falling on earlier non-target suppressed cells, and second, the increased strength and sparsity effect that can boost patterns that fall on earlier non-target excited cells.

In sum, experiments separating sequences in time (Fig. 4A-F), and constructing sequences designed to be excited or attenuated (Fig. 4G-K), together support a recurrent network mechanism for sequence tuning. Responses to an earlier pattern in a sequence create non-target responses in other local neurons that influence responses to later patterns.

## Discussion

Here we find that individual V1 neurons’ responses are modulated by inputs arriving to other V1 neurons earlier in time (Fig. 2). When we vary those prior inputs by changing prior visual stimuli, we find that earlier inputs boost responses to later inputs when the input sequence is derived from input sequences found in natural vision (Fig. 3). We show that a given flashed natural scene produces a response that lasts more than 100 ms (Fig. 3J), meaning the integration window for the interaction of one pattern with a future pattern can be at least this long. We then confirm that patterns interact across time via recurrent influences. To do this we design sequences of input patterns that should be boosted or attenuated if non-target responses to prior patterns interact with later patterns. We find that the resulting sequences are indeed selectively boosted or attenuated, as predicted. One intriguing result is that the boosted sequences produce sparser results than the attenuated sequences. This is consistent with the sparse responses to natural visual input observed by Gallant et al,^32^ and this observation supports the idea that recurrent interactions do shape responses to natural visual input.

Our results, taken together, show a computational role for the dense recurrent network in layer 2/3 of sensory (visual) cortex. While V1 does not generate complex sustained or ongoing dynamics, we find the V1 network does in fact support complex temporal processing, not by generating sequences but by filtering sequences of input.

It is likely that past work has not identified this sequence filtering phenomenon because physiological recording methods are limited by the curse of dimensionality. That is, it is impractical, in finite experimental time, to show sufficient stimuli to fully estimate models of receptive fields that encode complex features of natural vision. The other relevant constraint imposed by recording methods is the inability to easily determine which parts of neural responses to a stimulus come from feedforward input, and which from local, recurrent interactions. For these reasons, prior studies observed that natural responses were stronger and sparser on average than expected from simple receptive field combinations.^32,33^ But they could not identify that response gain varies depending on prior visual stimuli; this would have required either an impractically-detailed receptive field model, or another way to identify how a group of neurons was affected by different patterns of input to other local neurons. Here, two-photon stimulation allows us to match one specific population pattern of input to the visual context in which it occurs. In that way we can test whether a single pattern influences another, and the direction of its effect, without needing to identify interacting patterns via sampling large numbers of patterns.

### Relation to prior work: structure in spontaneous activity, straightening, cortical suppression, and hippocampal sequences

Prior observations that an echo of natural input can be seen in spontaneous activity^57–60^ are consistent with what we find. During spontaneous activity, neurons in the cortex fluctuate^61–63^ with measurable, but weak, correlation to natural visual responses. Our data suggest that while sequences of neural responses during spontaneous activity states are largely suppressed by the cortical network, some spontaneous patterns which partially match natural patterns can be weakly amplified by the recurrent network. However, because responses during spontaneous activity lack the structured feedforward driving inputs that occur during vision, spontaneous patterns are not amplified to the same extent as visual inputs, resulting in the relatively weak spontaneous correlations previously reported.

Martin and Douglas^3,4^ noted the large number of local recurrent synapses in the cortex and hypothesized that they could be used for amplification of input. Our work confirms that hypothesis, and extends their ideas by showing the kinds of inputs that can be amplified, including complex temporal statistics.

Another related observation is the straightening of perceptual and neural responses seen in natural vision^64,65^. Straightening refers to the relationship between response patterns at different moments of a natural movie. Prior work has found that cortical responses at one moment are more geometrically similar to later responses than is seen in the cortical inputs. The sequence filtering by the recurrent network we find could well be the mechanism for the straightening effect.

Stimulating one cell in visual cortex leads to an average suppressive effect that falls off with distance from the stimulated cell (also seen for small input patterns,^16,66^ confirmed for balanced networks in simulation^67^). This effect is also consistent with our findings. The suppressive effect seen in these studies is observed by averaging across many neurons at many distances, and appears to be dependent on the presence of visual stimulus drive,^67,68^ previously seen for wide-field stimulation as with one-photon optogenetics.

A final related prior observation is the pervasive suppressive response that occurs when external cortical stimulation ends, widely observed in electrical microstimulation studies.^69,70^ This effect is that stimulation — input drive applied to the cortex — is followed by an average suppression that lasts for approximately 150 ms. This duration is consistent with the timescale of suppression we find following a flashed natural scene (Fig. 3J). Our results put forth a possible explanation for the previously-reported average suppression: that external stimulation triggers the non-target effects that we find for input patterns (Fig. 4). However, using cell-specific stimulation we find the firing rate of individual non-target cells can be either higher or lower based on the previous input pattern (Fig. 2). The average suppression previously reported reflects the average of many cells whose target (electrically stimulated) or non-target status is not known. Two-photon optogenetics allows us to separate the two groups of neurons since we can identify the cells receiving input. The post-stimulation suppressive modulation previously reported thus confirms the non-target effects we report here.

Last, our sequence effects are fundamentally different than the sequential replay that is observed in the hippocampus.^71^ Our effect depends on firing rates over a few tens or hundreds of milliseconds, a distinct effect from the ordered spike sequences with millisecond-level precise timing that are generated during sharp-wave replay.^72,73^

### The dense E-E cortical network in sensory cortex may encode the structure of natural sensation

Our data suggest the large number of excitatory-excitatory recurrent connections in the cortex are used for learning the structure of the natural world. Depending on which cells are activated at a previous time, the recurrent network creates effects in other neurons that allow relative boosting (or suppression) of responses to particular input sequences.

The numerous excitatory recurrent connections in the cortex can, in principle, give the V1 network great capacity to selectively process the space of natural visual inputs.^36^ The patterns of population responses to visual input are high-dimensional: any given frame from a natural movie can affect the firing of many thousands of neurons in the cortex. So there is a large space of neural population responses, the set of patterns of activity to many different frames or different visual stimuli. While some sorts of dendritic nonlinearities (discussion in^74^) and nonlinearities due to short-term synaptic plasticity^75^ may also contribute, our data suggest that the high-dimensional space of visual input can be met with high-dimensional processing in the recurrent network. That is, the recurrent network has a large number of synapses which can act to set the non-target pattern produced by any input pattern. There are hundreds of millions of excitatory-excitatory recurrent synapses in a cubic millimeter of cortex,^1^ and so the dense recurrent network of the cortex seems well-placed to support the space of transformations required for sequence filtering.

While specific inhibitory connections^76,77^ may play a role in producing these non-target recurrent responses, no specific inhibitory population is needed to drive our effects. The suppression we observe could be generated via withdrawal of excitation. That is, the mechanisms we observe may be controlled primarily by excitatory-excitatory synapses, without being traceable to a set or subclass of inhibitory cells. In our experiments, we express opsin in excitatory cells only, so the interactions we observe must be caused by activation of excitatory cells. This excitatory activation can cause suppression in other excitatory cells in steady-state. For example, exciting excitatory cells may drive a few spikes in a number of inhibitory cells, which then increase their input to a large number of excitatory cells. In turn, the excitatory cells firing rates decrease, and that withdraws excitation from other connected excitatory cells. In this way the steady-state firing rates of the excitatory network can be primarily controlled by the pattern of connections amongst the excitatory cells, not by specific inhibitory connections, and without subsets of inhibitory cells that are substantially activated. That is an attractive principle for high-dimensional coding, since the space of excitatory-excitatory connections is larger than the space of inhibitory-excitatory connections, given the larger number of excitatory cells in the cortex. This type of mechanism, with excitatory activity in one population driving either excitation or suppression in other excitatory cells in the presence of balancing inhibition, is consistent with the balanced state or inhibition-stabilized state of cortical operation.^62,63,67,68,78,79^ Future work should be able to determine the role of excitatory and inhibitory interconnectivity in recurrent cortical computations.

The sequence filtering computation is a form of predictive processing.^27,80^ Because responses to a given visual input are influenced by previous inputs, the natural, expected sequence of inputs is boosted. This predictive processing — changing the response to the next input based on the previous — is achieved by altering the network’s input-output transformation, or network gain based, on prior inputs. This occurs via short-timescale non-target recurrent effects (Figs. 2,3), without large error or mismatch signals that might be expected from traditional predictive coding theories.^27,81–84^

### Sequence filtering seems likely to occur in other sensory regions beyond visual cortex

While we have made these measurements in visual cortex, we might speculate that recurrent connections in other sensory regions — auditory, somatosensory, etc — also are used for active filtering.^74^ Those sensory modalities also process natural inputs that vary in time, though often with faster changes than in vision, predicting different timescales of sequence filtering in other sensory regions. Additionally, associative brain areas downstream of primary sensory cortex can have tuning for complex sequences,^85^ and may filter at longer timescales.^86^

Beyond biology, recurrent networks in artificial systems also often are used to create temporally-structured computations. Our model findings (Fig. 3) are consistent with the idea that recurrent artificial networks can learn temporal statistical structure,^87^ as seen also in transformers like ChatGPT^88^ that have been used to generate highly complex natural language sequences. Thus, our results confirm the idea that densely-connected recurrent networks are useful for sequence processing and learning temporal structure, both in artificial systems and in biological brains.

## Conclusion

These data show a new and powerful purpose for recurrent connectivity in the sensory cerebral cortex: to confer sensitivity to sequential input. The visual cortical L2/3 network is sensitive to dynamic visual context, boosting responses to sequences of input corresponding to dynamic natural sensation, and suppressing others.

## Methods

### Animals

All experiments were conducted in adherence to NIH and Department of Health and Human Services (HHS) guidelines for animal research and were approved by the Institutional Animal Care and Use Committee (IACUC) at the relevant institutions (optogenetic experiments: NIMH; Neuropixels experiments: University of Washington.)

#### Optogenetic experiments

Emx1-Cre animals (N=11, https://www.jax.org/strain/005628)^89^ of both sexes were used. No systematic differences were observed between males and females. Viral injection and window implants were done at ages 2-7 months. After procedures, animals were singly housed on a reverse 12-hour dark/light cycle.

#### Neuropixels experiments

Male C57BL/6J mice (N=2), aged two to three months, were used. After procedures, animals were singly housed on a standard 12-hour dark/light cycle.

All animals were put on water schedule five or more days after head-plate surgery, and their weights were carefully monitored to ensure they remained above 85% of their baseline body weight.

### Viral injection and cranial window implants

We performed injections and implants as described in ^48^ (optogenetic experiments; NIMH) or ^90^ (2023; Neuropixels; University of Washington). Minor differences in the two sets of approaches are not expected to impact results.

#### Optogenetic experiments

Briefly, mice were given dexamethasone (3.2 mg/kg) 30 minutes before surgery and anesthetized with 1-3% isoflurane (in 100% O_2_). A titanium headplate was affixed using C&B Metabond (Parkell) and a 3 mm craniotomy was made over the primary visual cortex (-3.1 mm ML, +1.5 mm AP from lambda) and a glass optical window chronically inserted. Mice were injected with either a mixture of two AAV viruses to induce GCaMP and stChrimsonR into the cells (mixed virus injections), or a bicistronic virus expressing both GCaMP and stChrimsonR in each cell.^48^ For the mixed injection, the fluorescent calcium indicator, (AAV9-syn-jGCaMP8s; titers: 5.0-10x10^12^ genome copies (GC)/ml) and the soma-targeted opsin (AAV9-hsyn-DIO-stChrimsonR-mRuby2; titer: 3.0x10^12^ GC/ml) were mixed in phosphate-buffered saline. In the case of the bicistronic virus injections, we used AAV9-hsyn-DIO-jGCaMP8s-p2a-stChrimsonR (titers: 2.9-5.9x10^12^ GC/mL). For each mouse, 3 to 5 injections (100 nL/min, 200 µm depth) were made to cover a wider area of the cortex. A custom made light-blocking cap was fixed onto the implant to limit ambient light exposure and prevent debris from contacting the window. Experiments began three weeks after these procedures.

#### Neuropixel experiments

Briefly, mice were induced into anesthesia with 5% isoflurane, and maintained at 2-3%. Carprofen (5 mg/kg) and Lidocaine (2 mg/kg) were administered for analgesia. A titanium head plate and a 3D-printed chamber were affixed using Metabond. Carprofen was administered at 0.05 mg/ml in the water post-surgery for three days. Five days following recovery, mice were put on water restriction. After two days, they were habituated to head fixation for two days, during which they received up to thirty random 5µL sucrose water 10% rewards. Subsequently, animals were exposed to a set of 20 natural images presented over 1000 trials, followed by a 5µL sucrose water reward for five days. This step was performed to provide a control training condition for another experiment. None of the 20 images in this step were used as part of the experiment mentioned in this work. One or two days before recordings, a 2 x 2 mm craniotomy was performed over the primary visual cortex using a dental drill and stereotaxic techniques. The area was then covered with silicone gel. This process was conducted under anesthesia procedures used for the head plate implant, and Carprofen (5 mg/kg) was given for analgesia. A cap was placed on the chamber to prevent debris from getting into the area around the craniotomy.

### Retinotopic mapping

Before optogenetic experiments, we determined the location of V1 in the cranial window using a hemodynamic intrinsic imaging protocol previously described in ^91^. Briefly, small visual stimuli were presented while 530 nm light was delivered using a fiber-coupled LED (M350F2; Thorlabs, Newton NJ). Hemodynamic response was calculated as the change in reflectance of the cortical surface between the baseline period and a response window starting 3 ms after stimulus onset. Imaging was done on a Zeiss Discovery stereo microscope with a 1x widefield objective through a green long-pass emission filter, acquired at 2Hz. An average retinotopic map was fit to the cortical responses based on the centroids of the hemodynamic response for each stimulus location.

### Two-photon holographic imaging and stimulation

Two-photon imaging and stimulation procedures are described in detail in ^48^. Briefly, animals were awake and alert under a 16x water-immersion objective (Nikon; Tokyo, Japan; NA=0.8) manually positioned over the implanted optical window. Imaging was done with a custom-built microscope and controlled by ScanImage software in MATLAB (The Mathworks, Natick, MA). Calcium responses were measured ∼100-200 µm below the surface of the pia (L2/3 of V1) with an imaging field of view of 414 x 414 µm. Imaging was performed using 920 nm wavelength light at 15-20 mW and frames acquired at 30 Hz.

Holographic stimulation was performed using a femtosecond pulsed laser (1030 nm, Satsuma, Amplitude Laser, or 1040nm, Monaco, Coherent Inc.) A spatial light modulator (SLM) shaped the laser wavefront to create stimulation patterns (10 µm diameter disks; 30, 60, 90, or 120 ms; 8-16 mW/target, 500 kHz pulse rate). The radial point-spread function (PSF) of diffraction limited spots generated by the SLM was 9.4 µm and the axial PSF was 54 µm.The laser was gated on during horizontal flyback periods and off during the imaging pixel acquisition to allow for approximately simultaneous stimulation and imaging. Reported stimulation power is the average power over on and off periods (i.e. reduced by a factor of 0.3 from the laser power measured without this gating.)

Visual responses were measured in awake, head-fixed mice viewing flashed single natural image frames (40° circular mask with neutral gray background; 120 ms) or full natural movie stimuli (full-field, 2 seconds, 30Hz frame rate) presented on an LCD monitor. Mice were given small volumes of water on 20% of trials (3 µL).

### Two-photon data analysis

We performed motion correction using the CaImAn toolbox^92^ and cell segmentation using Suite2p.^93^ All data analysis was done in Python (https://www.python.org). For pixel based analyses, we computed ΔF/F_0_ (ΔF/F_0_; F: raw fluorescence intensity; F_0_: average fluorescence across the 45 imaging timepoints, 1.5 s, prior to stimulus presentation) at every pixel of the image stacks for display across the FOV. The ΔF/F_0_ for all time courses was calculated from fluorescent traces output from Suite2p. For full sequence trials (ABC, CBA), averaging windows for each pattern (A, B, C) started at stimulation onset of that pattern and lasted 300 ms; shifted for pattern order so the response window always began on the first frame of stimulation of that pattern (raw time course data without shift is shown in Fig. 2B). To isolate the optogenetic response following visual input, we subtracted off the response to the visual stimulus alone, [(vis+opto)-vis] (Fig. 3). Averaged response interval for individual cells began at optogenetic stimulus onset and ended at stimulus offset (120 ms, Fig. 3I). Activity of the C pattern in ABC sequence was calculated as [ABC - AB] (Fig. 4D,E). In Figure 4I-K, the averaging interval for individual cell responses started at stimulus onset and lasted 200 ms. Response intervals for all pixel based analyses started at stimulus onset and lasted 300 ms (Fig. 3B-E, 4A). Deconvolution (Fig. 1J) was done using OASIS with an autoregressive constant of 1.^94^ Population sparsity^32^ (Fig. 4K) was calculated as:

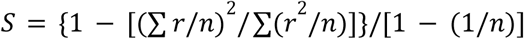

Where 𝑟 is the cell responses in the frame following the last stimulation and 𝑛 is the number of cells.

### Statistics

All statistical analyses were performed using Python. Statistical tests used include t-tests (Fig. 2D, Fig. 3I, Fig. 4D-F,I,K), Kolmogorov-Smirnov and Anderson-Darling tests (Fig. 2E,N-O, Fig. 4J), and Mood’s median test (Fig. 2Q). Significance threshold was held at 𝛼=0.05; n.s., not significant (p>0.05); *p≤0.05, **p≤0.01, ***p≤0.001. All experiments were replicated in multiple animals.

### Cell pattern selection

For analysis of visual response to flashed natural images, individual cell responses were averaged over a 300 ms interval starting from the stimulus onset. Cells activated by the visual stimulus were defined by responses exceeding a 5% ΔF/F_0_ threshold. All cells that met this threshold were included in the optogenetic pattern (11-15 cells per pattern, Supp. Fig. 2D,E).

To construct the boosted and attenuated patterns (Fig. 4), we first selected at random (not based on visual responses) the first 15-20 cell pattern in the sequence. Average ΔF/F_0_ response to stimulation of that pattern was calculated across a 300 ms time interval starting at stimulation onset. Excited non-target cells meeting a minimum threshold of 5% ΔF/F_0_ were included in the next pattern for the boosted sequence (15-20 cells per pattern). Conversely, suppressed non-target cell responses below -5% ΔF/F_0_ (15-20 cells per pattern) were selected for the attenuated sequence pattern. If more than 20 cells met these criteria, the 20 cells with the largest magnitude response were used. This process was repeated for the two sets of first and second patterns to generate the third pattern for each sequence. Each pattern in both sequences had an equal number of cells. Any cells included in previous patterns were excluded from selection for the subsequent patterns (Supp. Fig. 3G-J).

### Electrophysiology

For Neuropixels electrophysiology the mice were head-fixed for the recordings while seated in plastic support with forepaws on a wheel,^95^ with the ability to move body parts other than the head but no ability to locomote. The behavioral state of the animals was monitored using two Basler acA2440-75um cameras at 560x560 resolution. One camera focused on the eye area was used to monitor the pupil diameter, and the other focused on the face was used to detect licks and ensure the animal was not in distress. Animals were awake during the duration of the experiment. Recordings were conducted from all layers of the primary visual cortex using Neuropixel 2.4 probes^96^ with a sampling rate of 30 kHz. The recording sites were targeted based on the stereotaxic coordinates in the CCF Allen atlas using the Pinpoint system.^90^ The reference for the recordings was set at the tip of the electrode. Three recordings were performed: one from the first animal and two (on subsequent days) from the second animal. Spike sorting was executed using Kilosort 2.5 (https://github.com/MouseLand/Kilosort; RRID: SCR_016422)^96^ on each of the four shanks separately.

Visual stimuli were presented on three 60 Hz screens (LG LP097QX1), surrounding the mice and covering 270 x 70 degrees (azimuth x elevation) visual angle. Each trial featured a natural image from a set of two, sourced from the Allen Institute Brain Observatory, and repeated across all screens. These images were equiluminant and underwent histogram equalization for contrast consistency. Images were warped to appear rectilinear from the animals’ viewpoint. Each image was displayed 30 times for 50 ms. A reward of 5 µL of 10% sucrose water was administered at 1150 ms post-stimulus offset to maintain engagement. The trials were randomly interleaved with each other and with additional trials from another experiment, which also included natural image presentations and sucrose rewards. The inter-trial interval followed an exponential distribution with a minimum and mean of 2 and 2.6 seconds, respectively.

All electrophysiology analysis was done in Python. Baseline value computed in a 40 ms window ending 5 ms prior to stimulus presentation. Response value was calculated across a 50 ms interval beginning 5 ms after stimulus presentation. Excited cells are those with average response greater than baseline; suppressed cells those with response less than baseline.

### Modeling

We trained a recurrent neural network (RNN) consisting of N=500 units, whose input dynamics for the *i*-th neuron are given by:

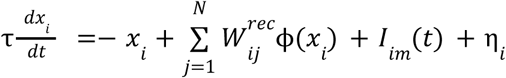

The input structure is defined by:

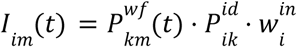

Where 𝑃^𝑤𝑓^is the input waveform, 𝑃^𝑖𝑑^ = {0, 1}, 𝑘 is the pattern, and 𝑚 is the sequence.

The readout of the network is defined as:

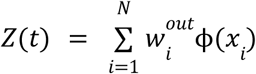

The transfer function of single units is ϕ(𝑥) = 𝑡𝑎𝑛ℎ(𝑥) . The weights of the input pattern 𝑤*_in_* are positive and exponentially distributed for a fraction 𝑝 = 0. 3 of units, and zero otherwise. The readout weights are an Identity matrix. The initial recurrent weights 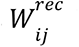, before any training, are independently sampled from a random Gaussian distribution with a mean zero and standard deviation 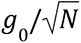. The noise term η_i_ is randomly sampled from a zero mean distribution with standard deviation 0.0005 at every time step.

We trained the recurrent weights 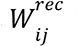 of the RNN using backpropagation-through-time (ADAM optimizer in PyTorch)^97,98^ such that the network readout 𝑍 for designated “trained sequences” matches a sparse scaled version of the time-varying input 𝐼_𝑖𝑚_(𝑡) where 30% of the cells have a target that is 2x their input and the other 70% have a target that is 0.5x their input. The network readout 𝑍 for “untrained sequences’’ was set to match the time-varying input without any scaling. Input structure consisted of 5 distinct patterns of units with independently sampled weights from the exponential distribution.

Sequences were generated by selecting from a set of 18 distinct input patterns. Untrained sequences were scrambled versions of the numerically ordered trained sequences. The input and output weights remained fixed. We trained the network on 500 epochs to produce these strong sparse responses. Control training had a target readout that exactly matched the time-varying input for all sequences. Parameters: τ = 60 𝑚𝑠, 𝑔_0_ = 0. 8, Euler integration timestep Δ𝑡 = 1 𝑚𝑠, learning rate 0.01.

---

**Supplementary Figure 1:**
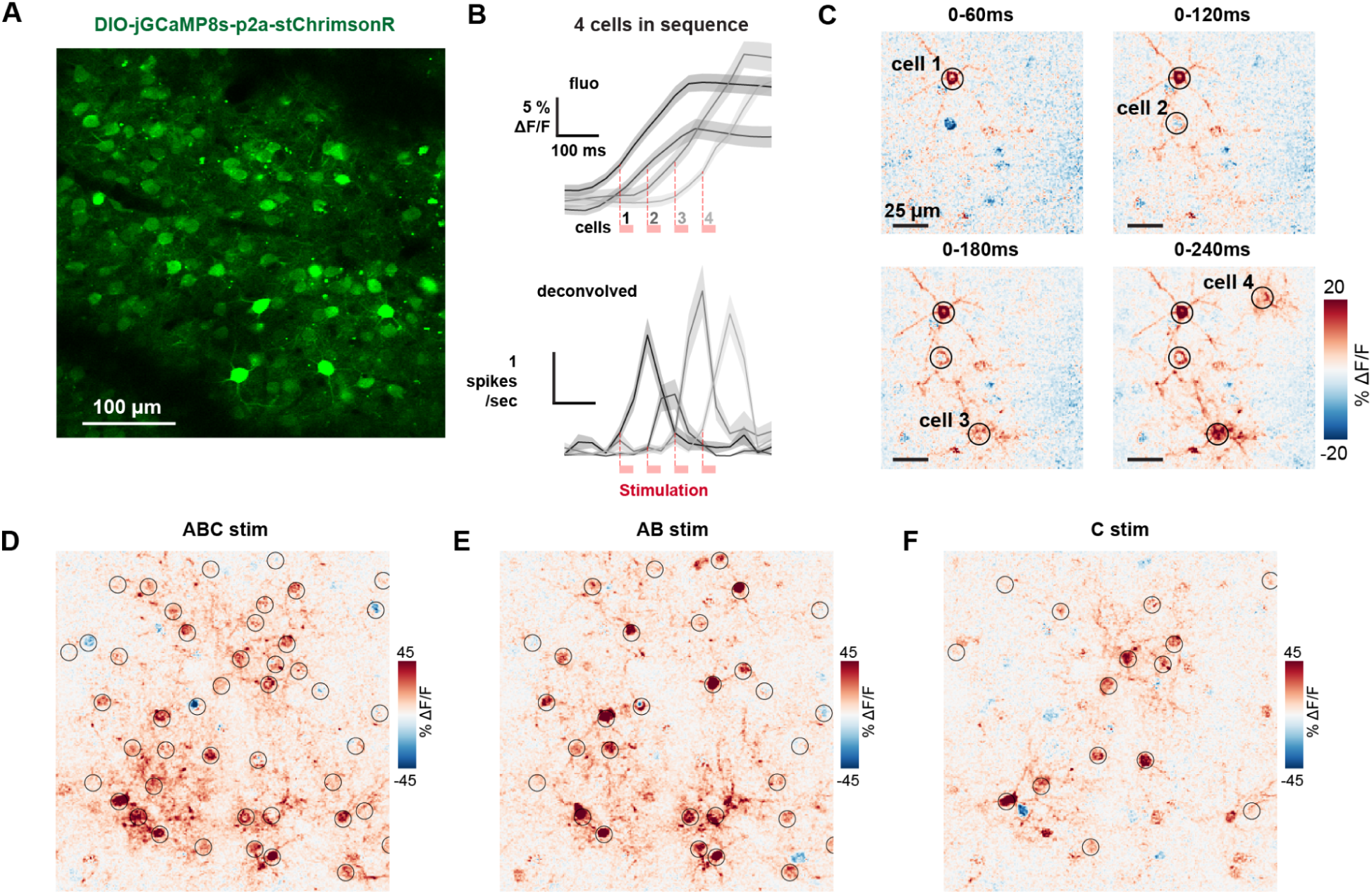
Two-photon stimulation responses: timing and individual patterns. **(A)** Example FOV image in L2/3 V1 with single virus injection, DIO-jGCaMP8s-p2a-stChrimsonR; this uses GCaMP8s which has even faster dynamics than GCaMP7s; more characterization in ^48^. **(B)** Sequential stimulation of 4 individual cells (30 ms per stim, 30 ms between stim). Fluorescent calcium traces above; deconvolved traces below. **(C)** Response maps from data in A. Stimulation pattern lasts 30 ms each in sequence triggered at times: 0 ms, 60 ms, 120 ms, 180 ms. b-c use preparation with expression of indicator and opsin in two viruses, AAV9-syn-jGCaMP7s and AAV9-syn-DIO-stChrimsonR. **(D-F)** Response maps of ABC, AB, and C pattern stimulation, using single bicistronic virus (jGCaMP8s, stChrimsonR, Methods). Associated with Figure 1 and 2.

**Supplementary Figure 2:**
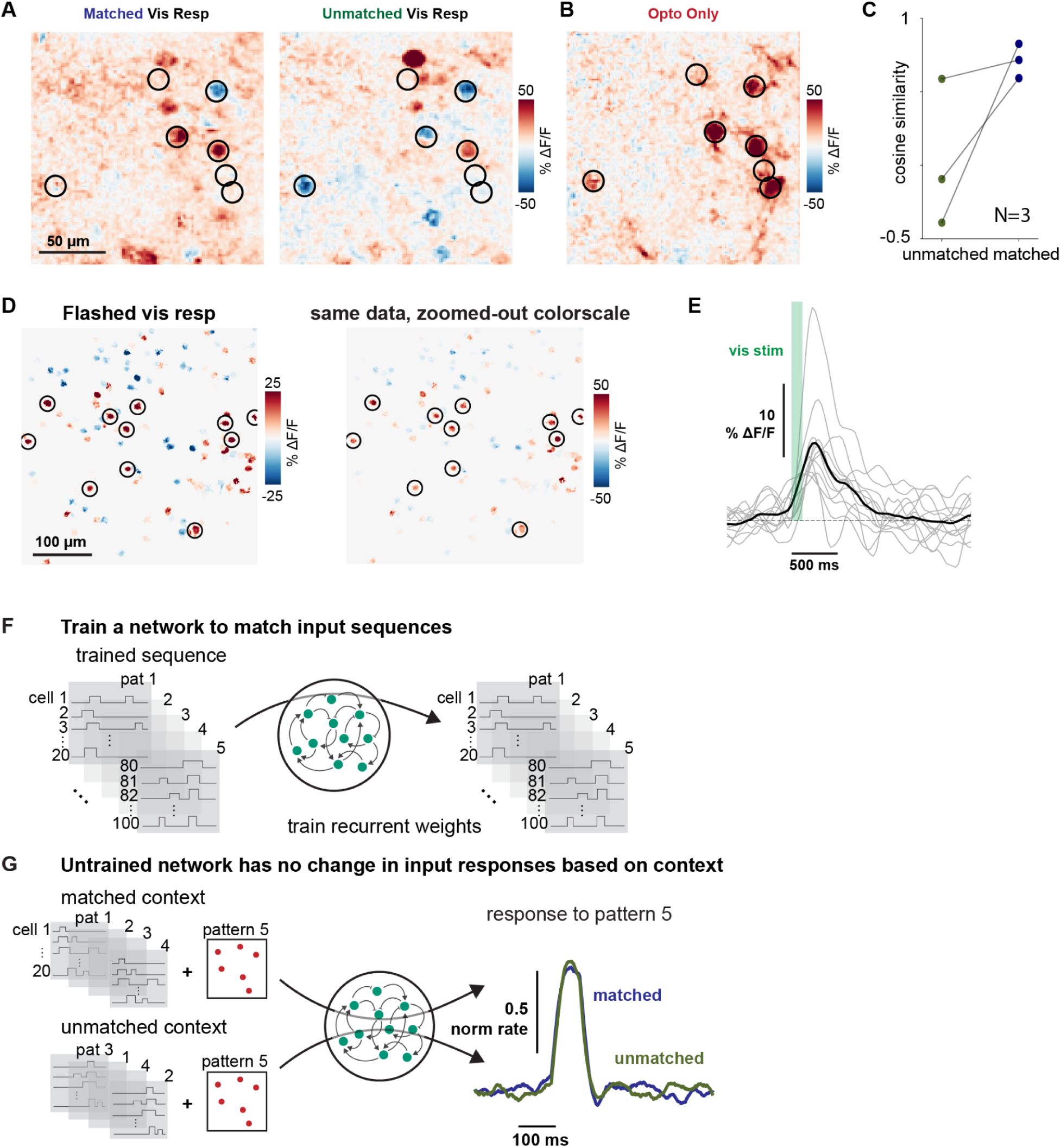
Boosting is not due to spike or indicator saturation. **(A)** Visual response alone of the same cells as shown in Fig. 3 for both matched and unmatched contexts at the time the optogenetic stim would be presented. **(B)** Optogenetic response of same cells without prior visual stimulus. **(C)** Greater cosine similarity between the optogenetic response alone and the visual responses alone at the time the optogenetic stim would be presented in the matched context than unmatched across experiments. **(D-E)** Cell selection for the natural-image-derived optogenetic pattern. **(D)** Response to flashed single frame; stimulated cells circled in black. Right: same data as left with un-zoomed color scale to show the larger cell responses that were chosen for optogenetic stimulation. **(E)** Timecourses of the circled cells in D. Only cells with a mean response above 5% are included in the pattern (Methods). **(F-G)** Control training for the artificial recurrent neural network. **(F)** Target output matched the input dynamics, no boosting. **(G)** Context dependent changes are eliminated when the model is not trained to selectively amplify sequences. Associated with Figure 3.

**Supplementary Figure 3:**
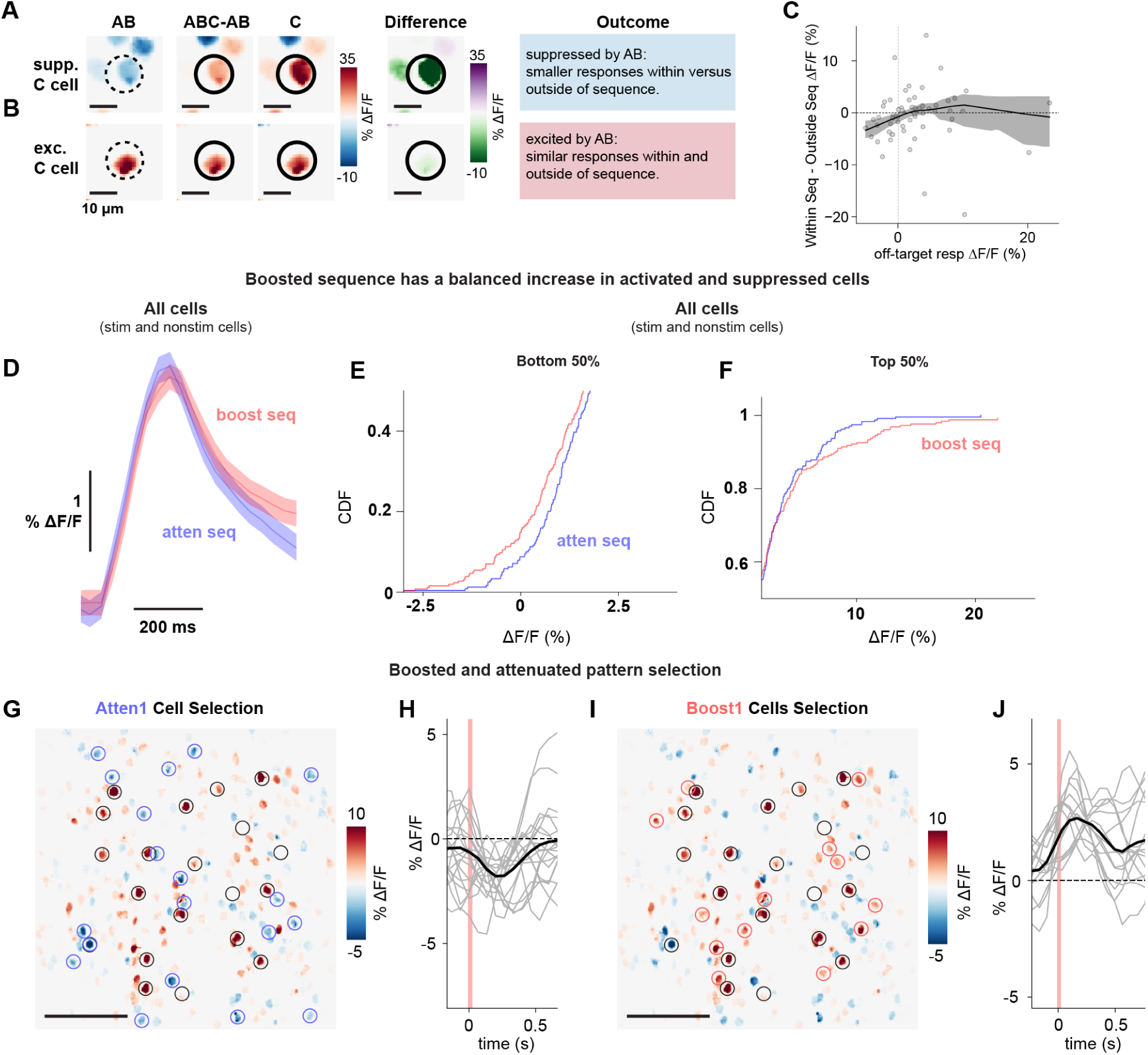
Constructed boosted and attenuated sequences have similar mean population responses. **(A)** Prior suppression attenuates the stimulated response. Example non-stimulated suppressed cell in response to AB stim. Dotted black circle: nonstimulated cell; Solid black circle: stimulated cell. **(B)** Prior excitation leaves the incremental response largely unchanged. Example non-stimulated excited cell in response to AB stim with same comparison within versus outside the sequence. **(C)** More suppression from prior stim leads to stronger attenuation of the cell response within the sequence. Lowess fit of C cell mean responses within vs. outside of the sequence ((ABC-AB) - C) depending on the non-target modulation from prior stimulation (AB). **(D)** While the stimulated cells show excitation/suppression (Fig. 4) as expected based on how they were selected, the overall population response mean does not appreciably change. Both stimulated and non-target (non-stimulated) cells. Average response (mean +/- SEM) of all cells in the FOV for the attenuated sequence (blue) and boosted sequence (pink). The mean population response for the boosted and attenuated sequences is similar. **(E-F)** Increased responses in the boosted sequence are balanced by an increased number of suppressed cell responses. Bottom 50% of cell responses for all cells in the FOV (D) for the attenuated sequence (blue) and boosted sequence (pink). Top 50% of cell responses (E). **(G-J)** Non-target patterns of suppression and excitation are selected for subsequent sequence patterns. **(G)** First arbitrary pattern (N=20 cells; pattern A) that was used to initiate sequence design circled in black. Selected suppressed (attenuated) cells circled in blue. This pattern contains cells that are suppressed by prior stimulation. **(H)** Timecourses of individual suppressed cells (from f) to A pattern stimulation. Mean trace (black) shows the mean is suppressed; suppressed cells were selected. **(I)** Same data as in C with boosted pattern cells shown. Boosted pattern: selected cells (red circles) excited by A stimulation. This pattern contains cells that are activated by prior stim. **(J)** Response timecourses of excited cells (F) in response to A pattern stimulation. Mean trace in black. Associated with Figure 4.

